# Temporal Dynamics of High-Frequency Oscillations in Alzheimer’s Disease: A Longitudinal Study in hAPP-J20 Mice

**DOI:** 10.64898/2025.12.10.693578

**Authors:** Keng-Ying Liao, Xu Han, Wen-Ying Chen, Wentai Liu, Yue-Loong Hsin

## Abstract

Alzheimer’s disease (AD) is characterized by progressive cognitive decline and increased seizure susceptibility; yet both the mechanistic and temporal links between AD and epileptogenesis remain poorly defined. In this study, we conducted a longitudinal analysis of epileptogenesis in relation to AD pathology using hAPP-J20 transgenic mice aged 9 to 27 weeks, encompassing the AD conversion phase. Wireless electroencephalography (EEG) was employed to monitor hippocampal high-frequency oscillations (HFOs), including ripples (80−200 Hz) and fast ripples (250−600 Hz), in conjunction with histological, behavioral, and neurophysiological assessments to characterize underlying neural circuit alterations. We identified three distinct epileptogenic stages in transgenic mice: (1) 9−15 weeks: emerging memory deficits, increased excitatory/inhibitory (E/I) neuron ratio, mossy fiber sprouting, and peak seizure-related mortality, alongside the initial emergence of pathological HFOs; (2) 15−21 weeks: a pronounced escalation of pathological HFO activity and persistent network hyperexcitability preceding detectable amyloid plaque deposition, indicating rapid epileptogenic progression; (3) 21−27 weeks: stabilization of HFO activity despite continued progression of amyloid accumulation, suggesting a plateau in epileptogenic remodeling amid advancing Alzheimer’s pathology. These findings indicate that neuronal hyperexcitability precedes amyloid plaque deposition and likely contributes to disease progression, highlighting a critical early window for therapeutic intervention in AD. Moreover, pathological HFOs hold promise as electrophysiological biomarkers of early circuit dysfunction and represent a promising target for modifying the course of AD.

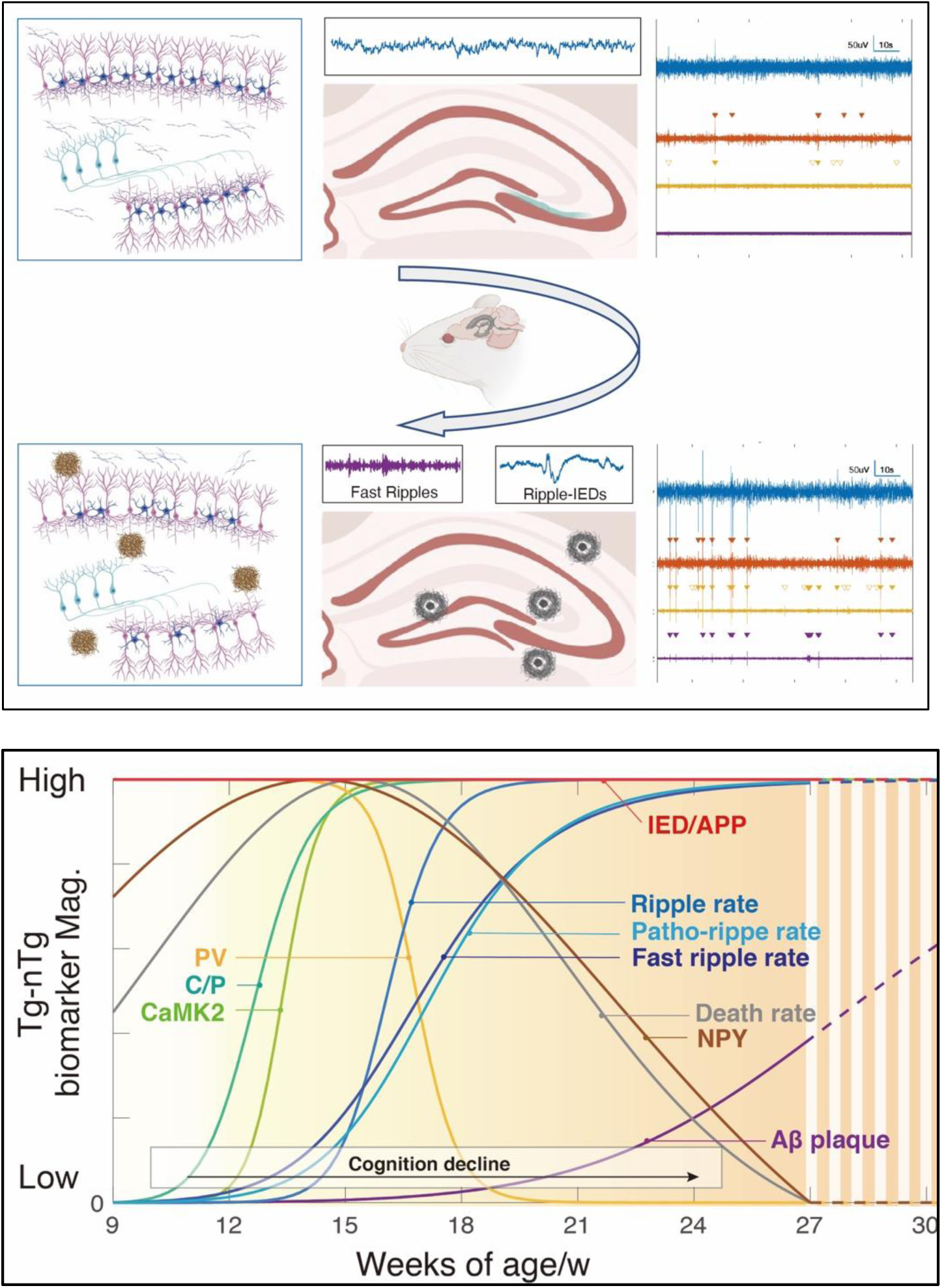

**Figure abstract:** This study examines the temporal dynamics of epileptogenic activity and amyloid pathology in hAPP-J20 transgenic mice, a model of Alzheimer’s disease (AD).

**Top:** Illustration of pre-plaque and post-plaque stages in hAPP-J20 mice, depicting neural network alterations, amyloid plaque deposition, and electrophysiological changes. The top panel shows the transition from a pre-plaque to a post-plaque state, where early hippocampal activity is stable and amyloid plaques are absent. As pathology advances, the network exhibits progressive hyperexcitability, characterized by increased pathological HFOs (ripple-IEDs, fast ripples), neuronal loss, NPY sprouting, and amyloid plaque formation.

**Bottom:** The progression of key biomarkers and disease hallmarks, comparing hAPP-J20 transgenic (Tg) mice with non-transgenic (nTg) controls. The bottom panel quantifies biomarker progression over time, revealing that CaMK2+/PV (C/P) imbalance emerges first, followed by a peak in pathological HFOs incident rate (patho-ripple, fast ripple rate), and finally, amyloid beta plaque (Aβ plaque). These findings establish a temporal relationship between early network hyperactivity and amyloid accumulation, underscoring the importance of targeting neural hyperexcitability as a therapeutic strategy to mitigate AD progression.

## 1. Introduction

Alzheimer’s disease (AD) affects over 55 million people globally and is the leading cause of neurocognitive decline. While the pathological hallmarks of AD—amyloid-beta (Aβ) plaques and neurofibrillary tangles (NFTs) of tau protein—have been extensively studied, a growing body of clinical evidence highlights the complex and dynamic relationship between AD and epilepsy. Approximately 10-42% of AD patients experience seizures, which are associated with worsened memory impairment and accelerated cognitive decline[1]. This association is particularly evident in late-onset epilepsy (onset after age 60), which may be recognized as a non-cognitive prodromal sign of AD[2, 3]. Despite this clinical overlap, the mechanistic and temporal interplay between epileptogenesis and AD pathology remains poorly understood.

Visible interictal epileptiform discharges (IEDs) have been associated with increased seizure risk and cognitive deterioration in AD [4, 5]. More recently, high-frequency oscillations (HFOs), brief oscillatory events in the 80–500 Hz range, have emerged as sensitive biomarkers for epileptogenic activity [6, 7]. While IEDs reflect meso-scale network activity involving broader neuronal populations, HFOs capture micro-scale network synchrony representing localized circuit dynamics. Abnormal HFOs indicate focal network hyperexcitability and have demonstrated predictive value for seizure onset; in epilepsy surgery, they are used to guide resection, leading to improved postoperative outcomes [8, 9]. Although HFOs have been extensively studied in epilepsy, recent observations of HFOs in AD animal models suggest that neuronal network dysfunction may be more intricately linked to AD pathology than previously recognized [9, 10]. These previous findings corroborate earlier reports of epileptiform activities in such models[11, 12] and underscore the potential role of neuronal hyperexcitability in driving memory impairment and cognitive decline.

Despite efforts to mitigate AD-related cognitive decline with anti-seizure medications, their efficacy has been limited. Although synaptic-modulating agents such as levetiracetam can reduce epileptiform activity in AD patients, these treatments have not demonstrated significant benefits in slowing or reversing cognitive deterioration[13]. This underscores the importance of precise timing in AD management, particularly concerning interventions aimed at preserving cognition. Given the progressive nature of AD, therapeutic strategies should be tailored to specific disease stages. Recent advances in Aβ-targeting monoclonal antibody therapies highlight the importance of early intervention, as their mechanism of action are most effective prior to extensive neurodegeneration. Likewise, the efficacy of anti-seizure or anti-epileptogenesis interventions may depend on their administration at precise stages of disease progression. However, the temporal dynamics of epileptogenesis in relation to amyloid pathology remains poorly understood, hindering the identification of optimal therapeutic timepoints.

Building on the foundational model proposed by Jack et al., which describes the sequential accumulation of amyloid and tau in relation to clinical and imaging markers, this study aims to address this critical gap. We hypothesize that HFOs serve as an early biomarker of neuronal hyperactivity associated with amyloid pathology, preceding conventional biochemical indicators of AD. Using hAPP-J20 transgenic mouse model, we longitudinally map electrophysiological and histopathological evidence of epileptogenesis, including neuronal population changes, mossy fiber sprouting, and HFO activity, alongside behavioral performance and amyloid deposition. By aligning these findings with established trajectories of human AD progression, this study seeks to elucidate the temporal dynamics between amyloid pathology and epileptogenic neuronal hyperactivity, ultimately providing a more refined framework for identifying therapeutic intervention windows in AD.

## 2. Methods

### 2.1 Animals and Experimental Design

hAPP-J20 transgenic mice (B6.Cg-Zbtb20^Tg(PDGFB-APPSwInd)20Lms^/2Mmjax) present a well-established mouse model for studying amyloid-based pathogenesis. This strain overexpresses human amyloid precursor protein (APP) and begins to develop stable amyloid plaques at approximately 5−7 months of age [14]. Other characteristic features, including neuronal loss, cognitive impairment, synaptic loss, alterations in local field potentials, have also been observed as early as 2−6 months of age. Therefore, this strain offers a valuable platform for investigating the early physiological effects of amyloid accumulation. Breeding pairs were obtained from The Jackson Laboratory (ME, USA) and subsequently bred and housed at BioLASCO (Yi-Lan, Taiwan). Mice were shipped to our institute at 7−8 weeks of age and maintained under a controlled 12-hour dark/light cycle environment with food and water *ad libitum*. Mice were acclimated to the facility for at least one week before being introduced to experimental procedures. All experimental procedures were approved by the Institutional Animal Care and Use Committee at National Chung Hsing University (IACUC No. 108-027) conducted in compliance with institutional guidelines. Transgenic mice were genotyped by standard polymerase chain reaction (PCR) following the genotyping protocol from the Jackson Laboratory (available at https://www.jax.org/Protocol?stockNumber=006293&protocolID=23978).

### 2.2 Behavioral Tests

The Morris water maze (MWM) is used to evaluate spatial learning memory and reference memory in mice. A circular pool (120 cm in diameter, 60 cm in height), marked with geometric visual cues on the inner wall to indicate four cardinal directions, was filled with tap water (25°C) and rendered opaque using non-toxic white paint. A complete MVM test spans a total of six days, consists of three phases: visible platform, hidden platform, and probe trial. On day 1, mice underwent visible platform test as a reference procedure to familiarize them with the task. In this phase, mice were trained to locate a 15 cm × 15 cm transparent platform submerged 1cm below the water surface and marked with a standing flag for guidance. From day 2 to 4 (hidden platform phase), the flag was removed, and the platform remained in a fixed location. Each mouse received four trials per day, starting from different quadrants in a randomized sequence. The cumulative distance moved and latency to reach the platform were recorded daily to assess learning performance. On day 6, mice underwent a 60-second probe trial during which the platform was removed. The cumulative time spent in the target quadrant and the frequency crossing over the former platform location were quantified to evaluate reference memory. The detailed procedure and protocol design were adapted from previously published study[15]All behavioral trials were captured and calculated through EthoVision XT video tracking system (Noldus, Wageningen, the Netherlands), with results verified through manual observation.

### 2.3 Immunohistochemistry

Mice were anesthetized with combination of Zoletil® (Tiletamine+Zolazepam, 20mg/kg) and Rompun® (xylazine, 5mg/kg) and exsanguinated via cardiac puncture (blood volume > 0.7 ml). Brains were excised and fixed in 10% neutral-buffered formalin for at least 24 hours, then embedded in paraffin and sectioned into 5µm slices. After deparaffinization, antigen retrieval was performed by immersing the sections in 0.1 M citrate buffer and heating in a water bath for 10 minutes, followed by enzymatic digestion with proteinase K (20 μg/mL in TBS) for 10 minutes. Blocking was performed using 3% hydrogen peroxide, washed with TBST, and incubated with the indicated primary antibodies, including APP (1:200, ab15272, Abcam, UK), Aβ42 (1:200, GTX134510, GeneTex, USA), CaMK2 (1:100, PA5-38239, ThermoFisher, USA), parvalbumin (1:400, MA5-35259, ThermoFisher, USA), neuropeptide Y (1:1000, ab221145, abcam, USA), diluted in 1% FBS. A biotinylated secondary antibody was applied following primary antibody incubation. After washing, the sections were treated with HRP-DAB for chromogenic detection and dried prior to mounting. Slides were scanned using the TissueFAX Plus system and the immunoreactive areas were quantified using HistoQuest single-cell analysis software (TissueGnostics, Vienna, Austria).

### 2.4 Hippocampal Electrophysiological Recordings

Mice were anesthetized with Zoletil® (Tiletamine+Zolazepam, 20mg/kg) and Rompun® (xylazine, 5mg/kg) for surgical implantation of a 2kHz high-sampling frequency wireless biopotential telemetry system (KAHA Science, now ADInstruments, Dunedin, New Zealand). Once the righting reflex and pain response disappeared, mice were secured in a stereotaxic frame, and two incisions were made: one at the back of the neck and one along the scalp midline. The transmitter was placed into a subcutaneous pocket made on the right lateral back. Recording electrodes were guided and implanted into the left CA1 region of the hippocampus CA1 (coordinates: ML: +1.2mm; AP: -2.0mm; DV: +1.5mm). The extracranial lead was fixed to the skull using dental glass ionomer cement. Incisions were closed with 5-0 nylon suture. Mice were given a single dose of ampicillin (200mg/kg) and Rimadyl® (carprofen, 5mg/kg) injection subcutaneously for postoperative care.

Hippocampal signals were recorded five days per week for up to four consecutive weeks post-implantation. Each recording session lasted a minimum of 10 minutes and was accompanied by synchronized video monitoring. As rodents are nocturnal, all recordings were collected during the light cycle. Only recordings obtained during sleep, defined as periods of immobility time longer than one minute, were used for further analysis.

### 2.5 Electrophysiological Signal Analysis

#### 2.5.1 Preprocessing Pipeline

The electrophysiological signal analysis process involved three steps: (1) pre-processing of intracranial EEG (iEEG) signals using a fine-tuned data-cleaning algorithm to remove artifacts and seizure activity, retaining only non-rapid eye movement (NREM) sleep segments; (2) detection of epileptic activities (EA), including IEDs and HFOs (ripples and fast ripples): (3) longitudinal analysis quantifying weekly incidence rate and band power of interictal EA in nTg and Tg mice from 9 to 27 weeks.

#### 2.5.2 Data Cleaning

A rigorous data cleaning protocol was implemented to achieve three primary objectives: (1) minimize false positives by excluding likely artifactual segments; (2) removing seizure segments to focus on interictal condition; and (3) retaining only segments from the NREM sleep stage, where both HFOs and IEDs are most frequently observed[7, 16].

For each recording, segments were excluded based on two criteria: a noisy high-frequency background, defined as an average background root mean square (RMS) value within a 5-second window in the 200–250 Hz band exceeding 1.7 µV (high-frequency threshold), or excessive background variability, characterized by a ratio of maximum to minimum background RMS values within a 5-second window greater than 10 (background variability threshold)[17]. The two thresholds were fine-tuned on a subset of recordings from the entire dataset (the fine-tuning sample size was estimated using statistical package from R). Each selected recording was manually evaluated and scored by two experts on a scale from one to four. Concurrently, a parametric sweep for the two thresholds was conducted on the data cleaning approach. The optimal threshold combination was determined by selecting the parameters that achieved the highest Cohen’s kappa coefficient between the averaged expert scores and the computer scores. Additionally, segments were classified as NREM sleep stage if their theta-delta ratio—calculated by dividing the average power in the 6–10 Hz band by the average power in the 2–4 Hz band—was less than two [18]. Only these NREM segments were retained for subsequent detection and analysis.

#### 2.5.3 Epileptic Event Detection

To detect IEDs from iEEG signals, a validated and effective detector based on a maximum likelihood estimation algorithm was applied to each cleaned segment[19]. The signal was first zero-phase bandpass filtered (10–60 Hz) and notch filtered (60 Hz) to reduce power line noise. The signal envelope was then calculated using the absolute value of the Hilbert transform and analyzed using a moving window approach (5 s, 80% overlap). For each window, the statistical distribution of the signal envelope was approximated using a log-normal model fitted via a maximum likelihood algorithm. Thresholds for IED detection were computed as three times the mode plus median of the fitted log-normal model. A threshold curve was generated by interpolating these values across windows using a cubic spline algorithm. Local maxima at the intersections of the signal envelope and the threshold curve were marked as putative IEDs, and events shorter than 120 ms were merged into a single IED.

To detect HFOs, including ripples and fast ripples, a validated detector was applied to each cleaned segment [20, 21]. The signal was zero-phase bandpass filtered in the 80–200 Hz range for ripple detection and 250–500 Hz for fast ripple detection, with 60 Hz notch filtering applied to reduce power line noise. The short line length (SLL) value of the filtered signal was calculated using a 5-ms sliding window. A signal was classified as a putative HFO if its SLL value exceeded 97.5th percentile of the empirical cumulative distribution function for each signal epoch. Additionally, events were required to contain at least six oscillations; events not meeting this criterion were excluded.

Following the detection of interictal epileptic activities, events where an IED co-occurred with both ripples and fast ripples were excluded from further analysis due to their high likelihood of being artifacts. Ripples co-occurring with IEDs (pathological ripples, or patho-ripples), were of particular interest, which are more strongly associated with pathological neural activity, such as epileptogenesis[22–24], compared to non-overlapping ripples.

Following the EA detection, both incidence rate and band power of EA were computed for each recording segment. The incidence rate was calculated by the number of detected EA events per second; band power ratio was calculated by the ratio of the EA band power (IED bandwidth: 10-80Hz; ripple bandwidth: 80-200Hz; fast ripple bandwidth: 250-500Hz) to the wideband power (<=1000Hz).

#### 2.5.4 Longitudinal Quantification

Longitudinal analysis was conducted separately for the Tg and nTg mice group. The incidence rates of epileptic activities (including IEDs, ripples, fast ripples, and patho-ripples), as well as the band power ratios of EAs (ripples and fast ripples), were averaged across all mice recordings on a weekly basis from weeks 9 to 27. To smooth the longitudinal incidence rate curves, a weighted linear least squares method with a first-degree polynomial model was applied. Linear regression was used to fit the band power ratio data.

Additionally, the incidence rates of EAs were averaged across all recordings for three developmental phases: early phase (9–15 weeks), middle phase (15–21 weeks), and late phase (21–27 weeks).

### 2.6 Statistical Analyses

For iEEG data analysis, given that within-subject variation was not significantly different from cross-subject variation, statistical comparisons between Tg and nTg groups were conducted weekly from week 9 to week 27 using Student’s t-test (*p* < 0.05). To evaluate group differences across developmental phases—early (9–15 weeks), middle (15–21 weeks), and late (21–27 weeks)—one-way analysis of variance (ANOVA) followed by Tukey-Kramer post-hoc analysis was performed, with significance set at p < 0.05. We computed pairwise similarity of residuals from a linear model fitted to the outcome variable without including subject-level random effects and found that within-subject and between-subject similarities were comparable ( Δ< 0.1 ), supporting the use of recordings as approximately independent in group-level analyses.

For other biomarker and behavioral data, statistical significance between the Tg and NTg groups were conducted every three weeks for the following measures: APP expression, Aβ plaque burden, mortality rate, neuropeptide Y (NPY), CaMK2, parvalbumin (PV) and the CaMK2/PV ratio. For MWM data, comparisons were performed monthly. Fisher’s exact test was applied for categorical variables (e.g., mortality rate). Student’s *t-test* was used for continuous variables (e.g., protein expression levels), with *p* < 0.05 considered statistically significant. One-way ANOVA with Tukey-Kramer post-hoc correction was used for phase comparisons (*p* < 0.05).

## 3. Results

### 3.1 Longitudinal Characterization of Amyloid Pathology and Early Behavioral Changes

In our study, hAPP-J20 mice, which carry transgene for human APP with Swedish (K670N/M671L) and Indiana (V717F) mutations, exhibited APP overexpression, promoting cleavage into aggregation-prone Aβ42 peptides. This process led to the progressive formation of Aβ plaques in the brain, initially depositing around 5−7 months and becoming pronounced by 8−10 months[25]. To delineate this temporal progression, we performed immunohistochemistry on mice sacrificed at three-week intervals (9-27 weeks), focusing on the hippocampus, to examine APP expression and Aβ plaque deposition (Figure 1A).

Most prior studies on Aβ plaque accumulation rely on cross-sectional designs, limiting resolution of the temporal dynamics. Although these studies suggest a sigmoid-like trajectory, they cannot directly validate this hypothesis without longitudinal tracking. In contrast, longitudinal amyloid PET imaging in human AD cohorts has confirmed this pattern, yet comparable evidence in animal models remains lacking. Our analyses revealed a progressive increase in Aβ plaque burden (quantified as Aβ42-positive area) starting at 18 weeks, with significant differences from nTg controls emerging at 21 weeks (*p*<0.05). By 27 weeks, plaque deposition had saturated, particularly emphasizing the 21−27-week interval as the initial phase of active plaque formation. Through a longitudinal immunohistochemical approach with closely spaced time points, our study captures the temporal progression of hippocampal Aβ accumulation in Tg mice. The trajectory follows a sigmoid-like pattern—a rapid early increase followed by a plateau—providing preclinical, time-resolved evidence that complements the dynamic biomarker model established in human Alzheimer’s disease. (Figure 1C, E, F, H, I).

**Figure 1.**
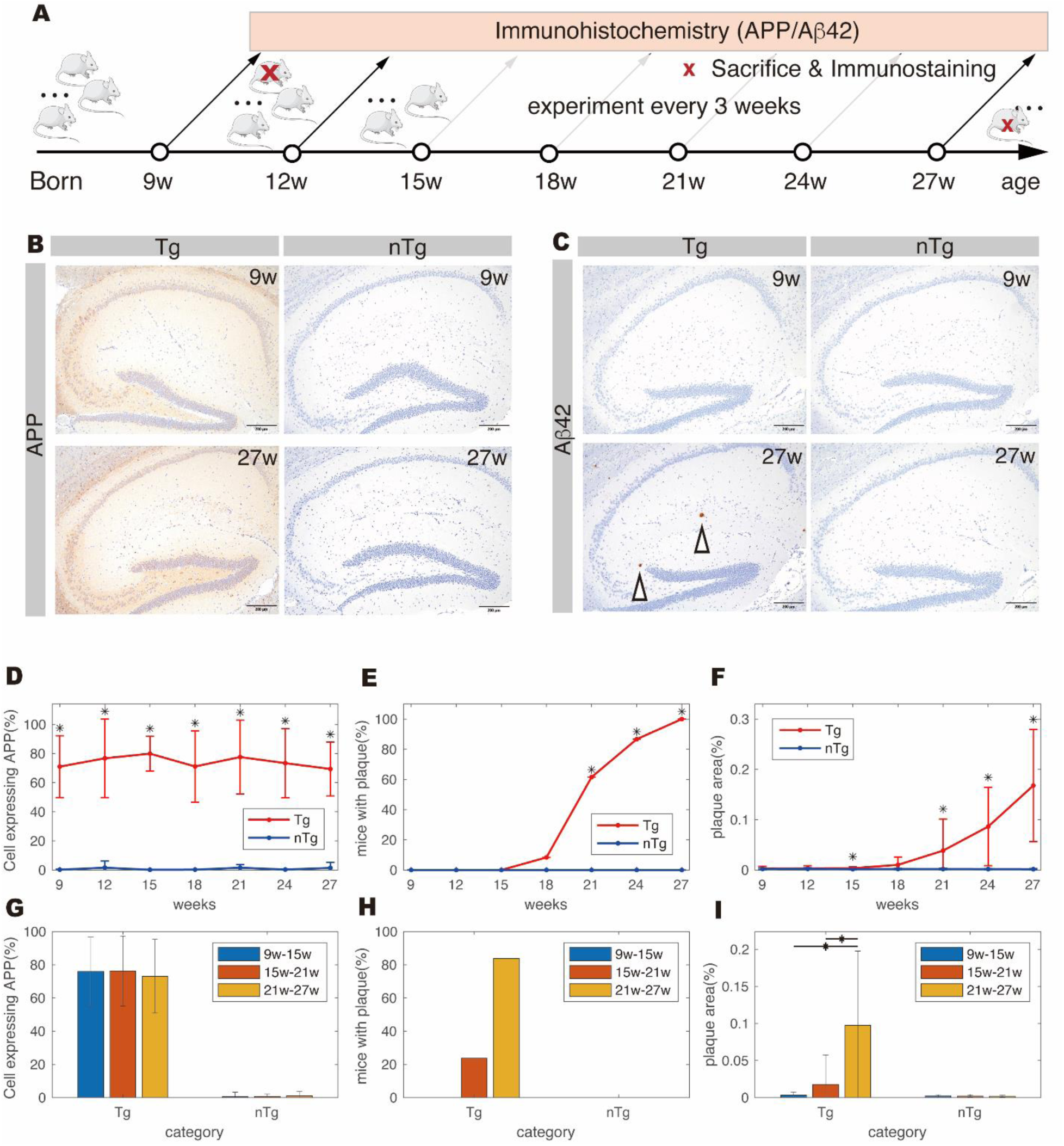
Temporal analysis of amyloid-related development. (A) Schematic illustration of the experimental timeline, including sacrifice and immunostaining for APP and Aβ42 conducted at three-week intervals. (B), (C) demonstrations of APP and Aβ42 immunostaining slides at 9 and 27 weeks of age. Arrows showing Aβ plaque deposition. (D), (E), (F) Line graphs showing the percentage of APP+ cells, the percentage of mice with plaque deposition, and the percentage of plaque-deposited area in Aβ42-stained sections across all experimental time points. (* p<0.05, n=8–10/per point) (G), (H), (I) bar chart comparing the percentage of APP+ cells, the percentage of mice with plaque deposition, and the percentage of plaque-deposited area in Aβ42-stained sections across distinct developmental phases (categorized as 9–15 weeks, 15–21 weeks, and 21–27 weeks). Data are presented as mean ± SD. (* p<0.05, n=26–29/per bar)

Despite the dynamic changes in plaque deposition, APP expression remained constant throughout the life of Tg mice. Notably, Tg mice exhibited significantly higher APP expression compared to nTg mice, emphasizing its dual role in the central nervous system (CNS) (Figure 1B, D, G). APP is a versatile transmembrane protein with critical roles in CNS function, including neuronal growth, synaptic formation, cell adhesion, and signal transduction. However, overexpressing APP in Tg mice leads to increased aberrant cleavage of Aβ, facilitating amyloid plaque formation and the early onset of Alzheimer’s pathology.

While amyloid pathology becomes apparent around 21 weeks of age, J20 mice have been shown to exhibit memory deficits in variant behavioral tasks at earlier age[26]. We evaluated learning and working memory using Morris water maze test in Tg and nTg mice, with assessments conducted every four weeks from 8 to 24 weeks of age (Figure 2A). Significant performance deficit in Tg mice was observed starting at 12 weeks of age, preceding the onset of notable Aβ plaque deposition (Figure 2B, C, D). This performance deficit remained consistent at 16, 20, 24-weeks of age but did not show gradient deteriorate with amyloid pathology progression. However, the time and distance spent reaching platform during the four-day Morris water maze test representing the learning curve gradually flattened along age. These results suggest that cognitive impairments in AD may arise from early molecular changes independent of visible plaque accumulation.

**Figure 2.**
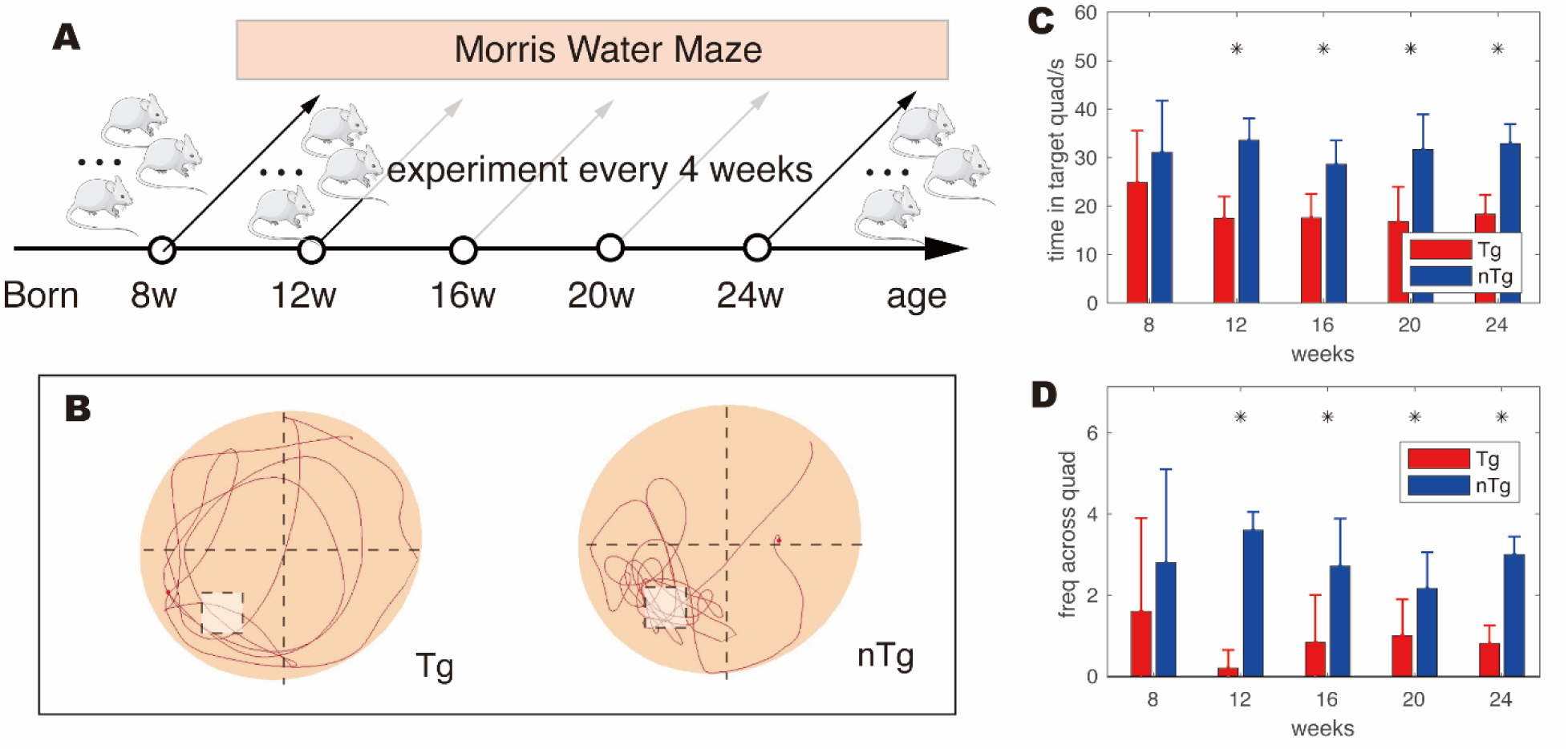
Temporal analysis of cognition in J20 mice by Morris water maze. (A) Schematic illustration of the experimental timeline. Morris water maze tests were conducted at four-week intervals from 8 to 24 weeks to assess spatial learning and memory. (B) Representative swim trajectories during one probe trial illustrate impaired spatial search in a single Tg mouse compared to a single nTg mouse. (C) Quantification of time spent in the target quadrant during the probe trial across all tested ages. Tg mice showed significantly reduced time in the target quadrant from 12 weeks onward, indicating early impairment in spatial memory. (D) Frequency of platform area crossings during the probe trial. Tg mice exhibited significantly fewer target crossings at all time points after 8 weeks, suggesting persistent deficits in spatial reference memory. Data are presented as mean ± SD. (* p<0.05, n=5/per bar)

### 3.2 Pre-Plaque Hyperexcitability and Hippocampal Circuit Remodeling

We observed convulsive seizures in J20 mice as early as two weeks of age (supplementary video), confirmed by intracranial hippocampal EEG (Figure 3F). Deceased mice often exhibited opisthotonos, indicative of heightened hippocampal excitability (Figure 3D). These deaths were not linked to external factors such as fighting, self-mutilation, or malnutrition. Tracking mortality at three-week intervals revealed an exponential increase, peaking at 15 weeks, before gradually returning to baseline levels (Figure 3B). This temporal pattern provides a critical window for studying early epileptogenesis.

**Figure 3.**
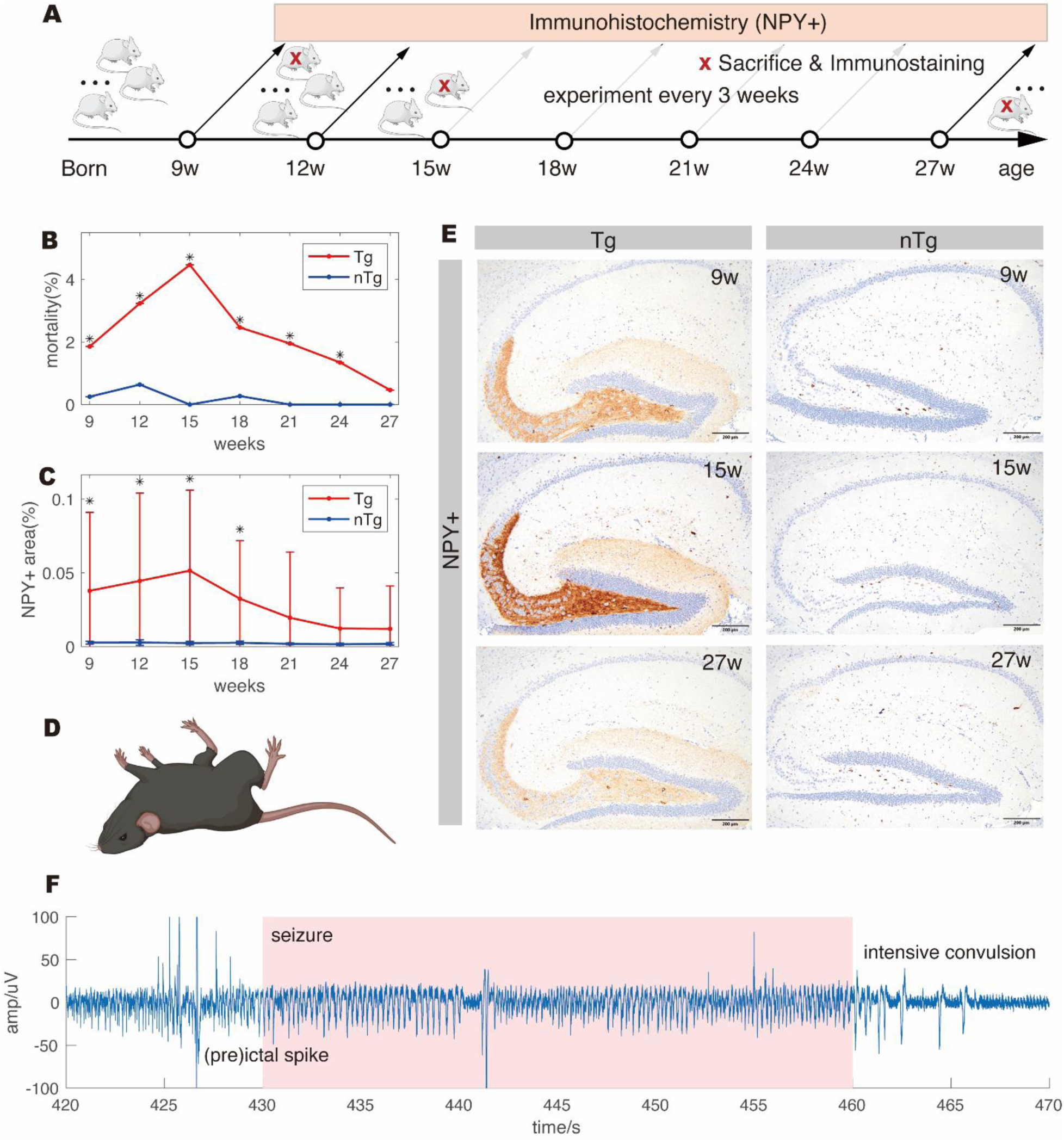
Remodeling of hippocampus circuits causing life threatening convulsions at the pre-plaque stage. (A) Schematic illustration of the experimental timeline, including sacrifice and immunostaining for NPY conducted at three-week intervals. (B) line graph showing the observed mortality rate calculated by a three-week interval across all experimental ages. A marked increase in mortality peaks at 15 weeks of age before gradually returning to baseline levels. (based on observation record of 1059 for nTg, and 730 for Tg) (C) line graph showing the percentage of NPY+ area in the hippocampus across all experimental ages. A strong correlation coefficient (r=0.899) was found between the percentage of NPY+ area and mortality rate. (* p<0.05, n=8-10/per point) (D) Illustration of convulsive seizures in mice, characterized by loss of balance and tonic-clonic convulsions involving the entire body. Mice that died of seizures were often found with stiff, straightened limbs. (F) Representative intracranial EEG recorded during a seizure episode. Data are presented as mean ± SD.

As mentioned earlier, APP and its proteolytic fragments provide various physiologic bioactivities. However, aberrant processing of APP could cause pathological effects and thereby alters neuronal excitability, which likely explains the higher mortality rate observed in Tg mice as early as 9 weeks of age. Interestingly, the mortality rate peaking at 15 weeks of age precedes the deposition of Aβ plaques in the hippocampus, suggesting an unrecognized dynamic under the formation of the amyloid cascade, particularly during soluble oligomers to insoluble fibrils and plaques. These observations underscore the detrimental impact of amyloid fibril processing, a prerequisite for amyloid plaque formation, contributing to the development of neural hyperactivity and life-threatening convulsions.

NPY, a neurotransmitter peptide widely distributed in the CNS, is predominantly expressed in the interneurons where it modulates glutamate release during high-frequency activity. Elevated NPY levels are transiently observed following acute seizures, suggesting a compensatory mechanism to reduce hyperexcitability[27]. In particular, studies of temporal lobe epilepsy (TLE) have reported increased NPY expression alongside characteristic neuropathological changes such as neuronal loss in the CA1, CA3, and hilus, mossy fiber sprouting, and gliosis. Reflecting this pattern, we evaluated NPY expressing cells in Tg and nTg mice, with assessments conducted every three weeks from 8 to 24 weeks of age (Figure 3A), and our longitudinal analysis revealed a significant increase in NPY levels in the CA3 and dentate gyrus of Tg mice (Figure 3C). These levels peaked at 15 weeks before gradually declining to baseline (Figure 3B, C, E), mirroring mortality dynamics with a strong coefficient correlation (r = 0.899). This parallel increase in NPY and mortality peaking at 15 weeks of age implied a temporary excessive seizure activity prior to amyloid plaque deposition, which curiously subsides as the disease progresses. However, the interaction between early neuronal hyperactivity and amyloid pathology, particularly its influence on amyloid fibril processing, warrants further detailed investigation to elucidate their causal relationship.

### 3.3 Alterations in PV Interneurons Reveal Emerging Excitatory/Inhibitory Imbalance

Pyramidal cells are the principle excitatory neurons in the hippocampus. These neurons, characterizing by their excitatory nature, synaptic plasticity, and complex firing pattern, forms a meticulous and networked circuit which is thought to be involved in memory encoding and retrieval [28]. PV-expressing inhibitory interneurons play a crucial role in regulating pyramidal cells due to their fast-spiking nature. They can generate rapid action potentials at highly synchronous frequencies, enabling them to precisely control the timing of pyramidal cell firing. Yet, these interneurons are highly sensitive to cellular stress and developmental stage-specific expression, makes them particularly vulnerable under diseased conditions, contributing to unstable neural circuits. Together, the dynamic interplay between these excitatory and inhibitory neurons is crucial for maintaining proper brain function. Therefore, we evaluated number of PV-expressing interneurons and CaMK2 pyramidal cells in Tg and nTg mice, with assessments conducted every three weeks from 9 to 27 weeks of age (Figure 3A),

Given the physical characteristics and critical roles of these neurons in the hippocampus, in our longitudinal observation, we examined counts of PV-expressing interneurons (labeled by PV) and pyramidal cells (labeled by CaMK2) in the hippocampus of J20 mice. We observed a progressive decline in the number of PV-expressing interneurons in Tg mice compared to nTg controls, with this reduction becoming more pronounced from 18 to 27 weeks of age (Figure 4B, E, H). While the number of pyramidal cells did not differ significantly from nTg mice, a slight increase was noted over time (figure 5C, D, G).

**Figure 4.**
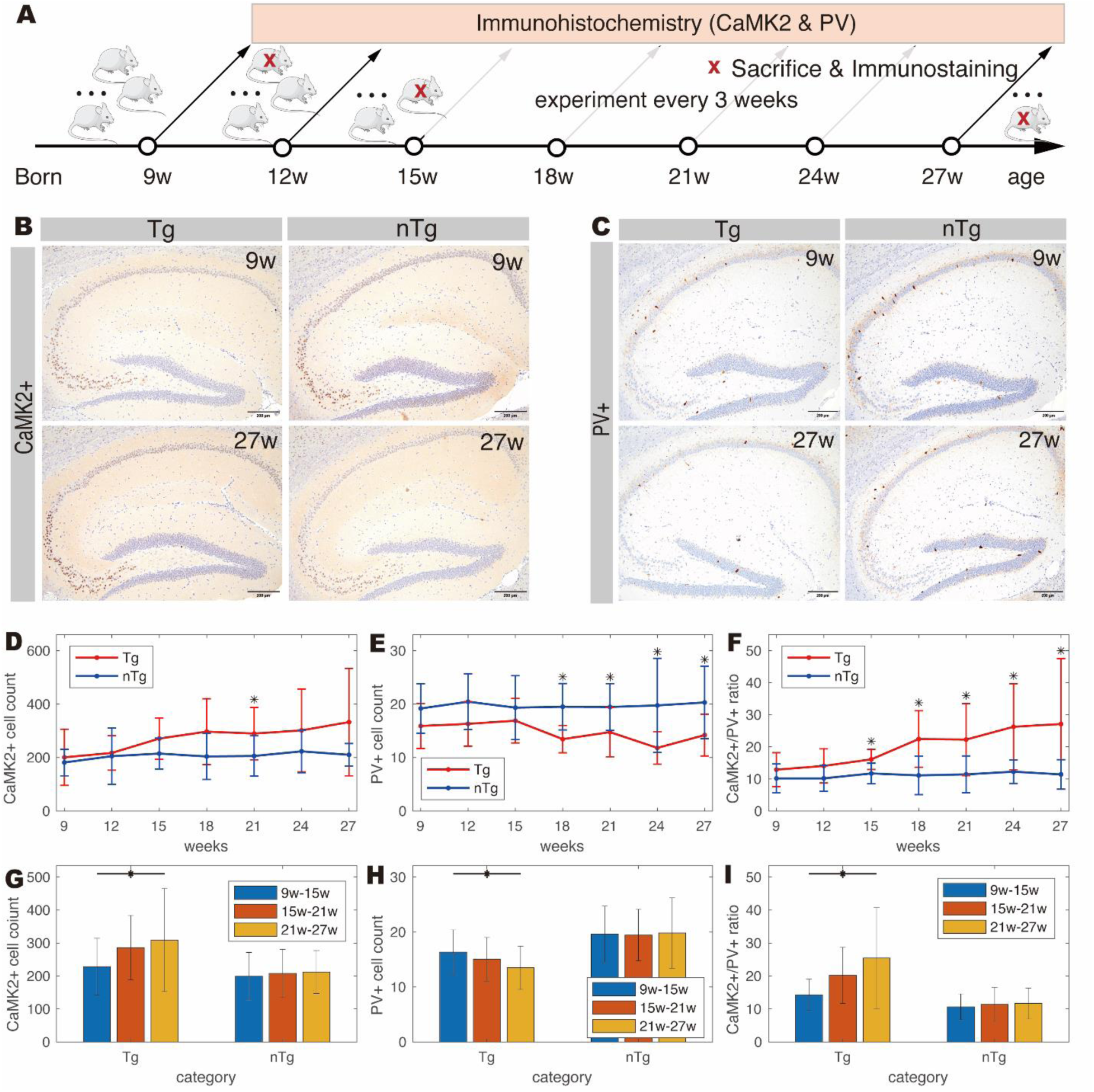
Alterations in excitatory and inhibitory neuron populations in J20 mice. (A) Schematic illustration of the experimental timeline, including sacrifice and immunostaining for CaMK2 (excitatory marker) and PV (inhibitory marker) performed at three-week intervals. (B), (C) Representative immunostaining images for CaMK2 and PV at 9 and 27 weeks of age. (D), (E), (F) line graphs showing the number of CaMK2+ cells, PV+ cells, and the percentage of CaMK2/PV across all experimental ages. (* p<0.05, n=8–10/per point) (G), (H), (I) Bar charts comparing the number of CaMK2+ cells, PV+ cells, and the percentage of CaMK2/PV across distinct developmental stages (categorized as 9–15 weeks, 15–21 weeks, and 21–27 weeks). Data are presented as mean ± SD. (* p<0.05, n=26-29/per bar)

**Figure 5.**
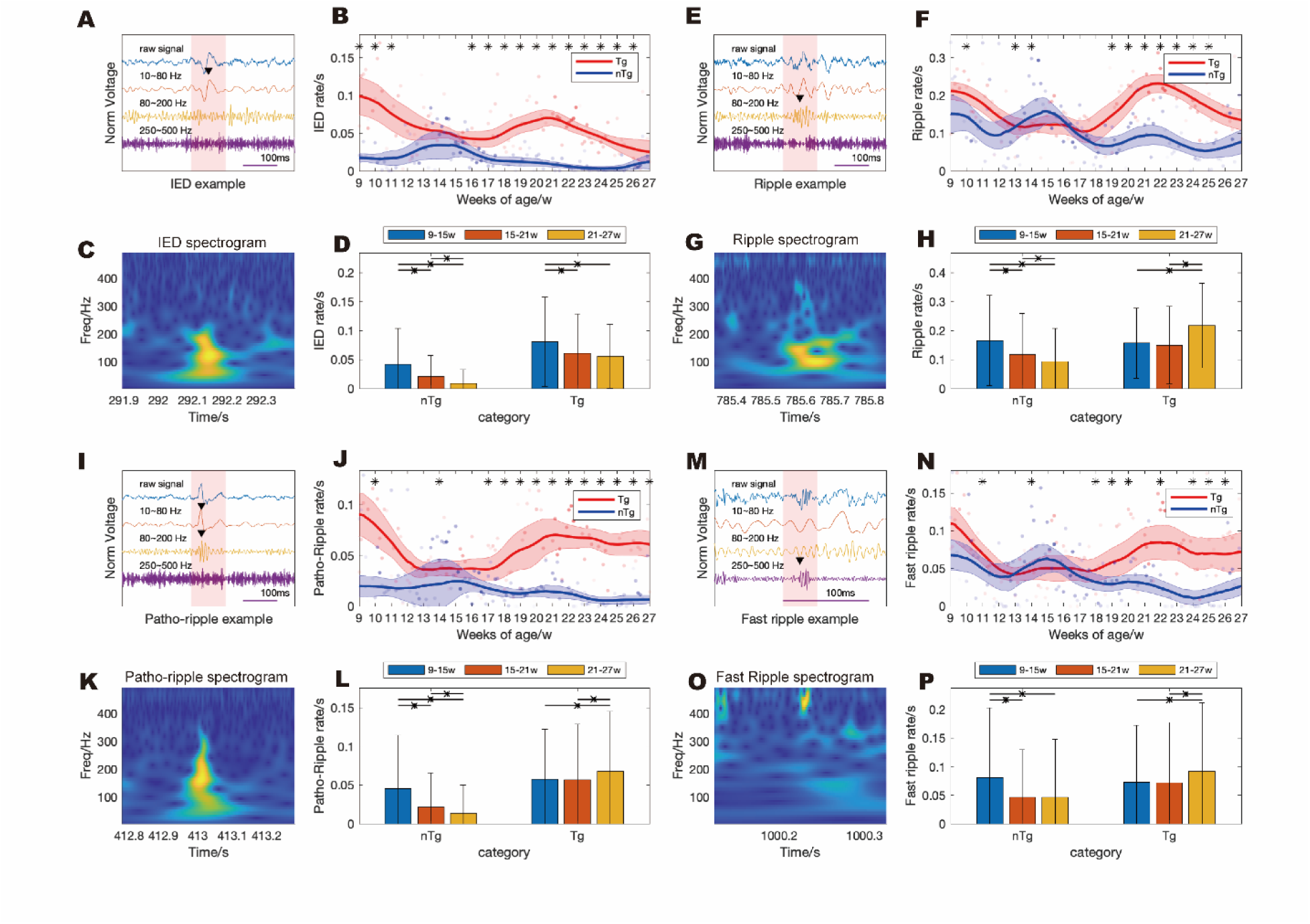
Experimental setting for chronic CA1 mono-electrode recording and temporal analysis of band power ratios in intracranial electroencephalographic (iEEG) activities. (A) Schematic illustration of the electrode implantation and daily iEEG recordings, continued until either the subject’s death or electrode detachment. (B) Demonstration of MT10B KAHA mono-electrode telemeters dimension along with implantation details. (C), (D) Temporal changes in ripple band and fast ripple band power ratios associated with EAs in Tg and nTg mice. Linear regression lines represent trends in band power ratios with age, and the shading indicates the 95% confidence interval (CI).

Notably, the ratio of pyramidal cells to PV-expressing interneurons in Tg mice increased significantly from 15 weeks and continued to widen as the mice aged (Figure 5F & I). This increased ratio of pyramidal cells to PV-expressing interneurons suggests heightened relative excitability in the brain alongside diminished inhibitory control. The resulting shift in excitatory-inhibitory balance lowers the seizure threshold and raises the likelihood of excessive, synchronized neuronal activity, thereby making the brain more susceptible to seizures, which partially explains the heightened seizure-related mortality observed earlier in J20 mice. These findings, in the context of amyloid pathology in J20 mice, provide insight into how early PV-interneuron depletion may contribute to hyperexcitability and epileptogenesis, paralleling mechanisms observed in other neurological conditions such as epilepsy.

### 3.4 Temporal Dynamics of Epileptiform Oscillations in the Hippocampus

Utilizing high-sampling-rate chronic CA1 monopolar electrode recordings (Figure 5B), we collected extensive intracranial EEG recordings spanning from 9 to 27 weeks of age, encompassing the pre-plaque to post-plaque deposition stages (Figure 5A). This approach allowed us to correlate hippocampal histopathological changes with seizure-related electrical activity.

In addition to analyzing visibly recognizable IEDs, such as sharp and spike waves, we examined HFOs, a biomarker of epileptic activity. HFOs are typically categorized into two frequency bands: low-frequency ripples (80−200 Hz) and high-frequency fast ripples (250−500 Hz). While ripples are mostly reflected normal physiological functions such as synaptic plasticity or memory consolidation, fast ripples are aberrant hyper-synchronous microcircuits strongly associate with epileptogenesis. They can appear ipsilateral in the seizure onset zone despite of the coupling effect, which reinforces their clinical value as a specific biomarker for localizing epilepsy. This characteristic makes them highly relied upon in epilepsy surgery for identifying the epileptogenic focus. In contrast, pathological ripples, which is ripple co-occur with IEDs, arises from network dysfunction often seen in regions where physiological ripples typically occur. Although pathological ripples are less region-specific than fast ripples, they provide valuable insights into neuronal network dysfunction and epileptogenesis[7]. Together, both pathological ripples and fast ripples—collectively known as pathological HFOs—serve as critical indicators for understanding local neuronal activity and epileptogenesis.

In our study, IEDs remained significantly elevated in J20 mice throughout the recording period compared to nTg mice, paralleling with the constant APP overexpression (Figure 6A−D). Ripple rates were comparable between nTg and Tg groups before 15 weeks of age (pre-plaque stage) but increased significantly in Tg mice after 19 weeks of age (Figure 6E−H)., coinciding with the emergence of patho-ripples (Figure 6I−L). Similarly, fast ripples rates progressively rose, becoming significantly elevated in Tg mice after 19 weeks of age alongside amyloid plaque deposition (Figure 6M−P). The band power ratio of HFOs steadily increased from week 9 to week 27 (Figure 5C&D) reflects enhanced strength in synchronous neuronal firing, indicative of heightened network excitability.

**Figure 6.**
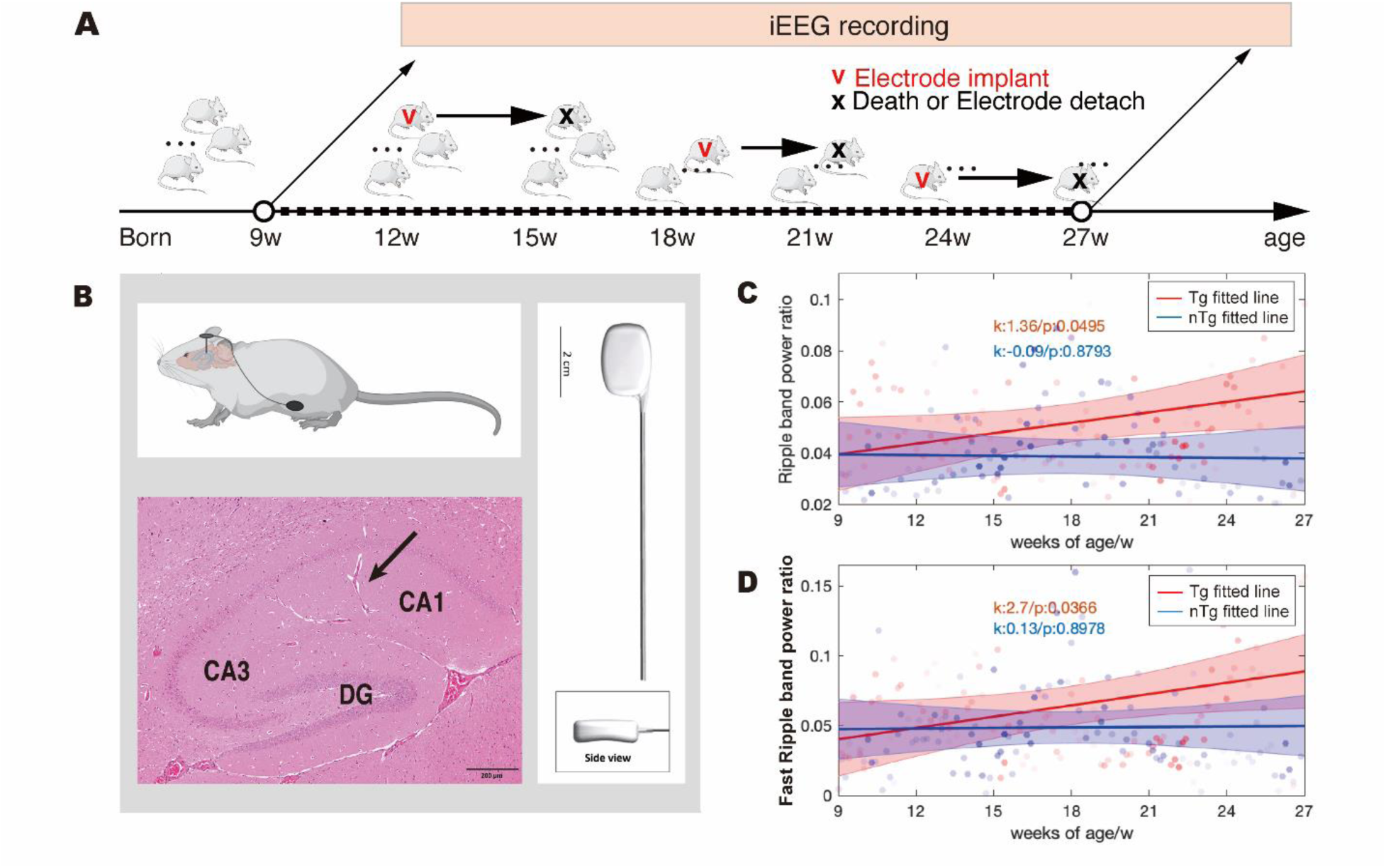
Temporal analysis of epileptic activities (EAs) incidence rates by chronic CA1 mono-electrode recordings. (A), (C), (I), (K) Representative waveforms of interictal epileptiform discharges (IEDs), ripples, patho-ripples, and fast ripples, respectively. (B), (D), (J), (L) Line graphs showing age-related changes in the incidence rates of IEDs, ripples, patho-ripples, and fast ripples in Tg and nTg mice. Shading represents the 95% confidence interval (CI). (E), (G), (M), (O) Spectrogram illustrating examples of IED, ripple, patho-ripple and fast ripple respectively. (F), (H), (N), (P) bar charts comparing the incidence rates of EAs (IEDs, ripples, patho-ripples, and fast ripples) across developmental stages (categorized as 9–15 weeks, 15–21 weeks, and 21–27 weeks). Data are presented as mean ± SD. (* p<0.05, n=33 for Tg, n=31 for nTg).

**Figure 7.**
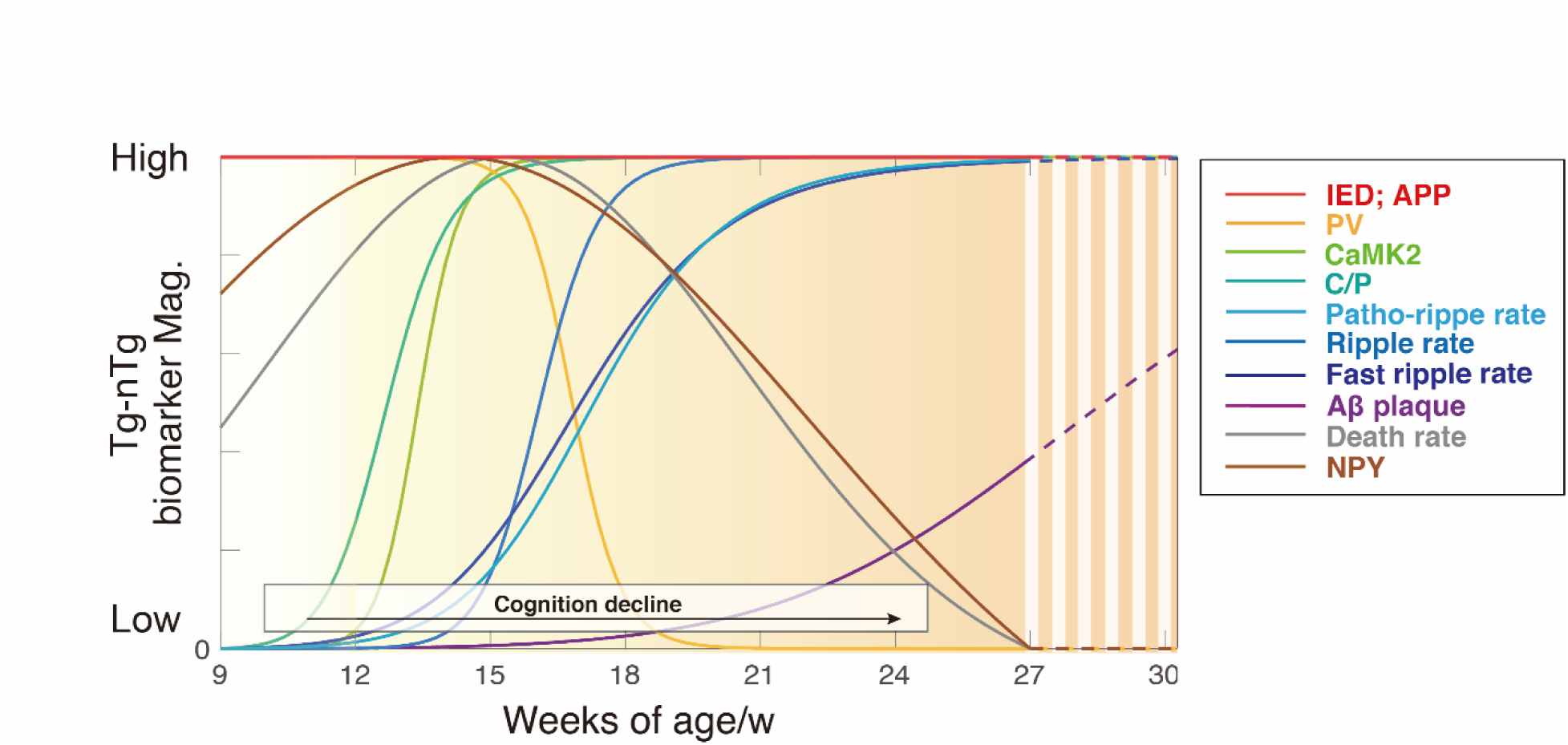
Various hallmarks change with age for the difference between Tg mice and nTg mice. The curves reveal that CaMK2+/PV (C/P) imbalance emerges first, followed by a peak in HFOs incidence rate (patho-ripple, ripple and fast ripple rate), and finally, amyloid deposition (Aβ plaque). These findings establish a temporal relationship between early network hyperactivity and amyloid accumulation, underscoring the importance of targeting neural hyperexcitability as a therapeutic strategy to mitigate AD progression.

In summary, the observed increase in HFO band power, followed by a rise in ripple and fast ripple rates, indicates a heightened state of neural activity characterized by increased amplitude and frequency of synchronized neuronal firing. This heightened activity reflects underlying changes in neuronal circuits during the amyloid pathology progression, such as enhanced synaptic plasticity, increased excitability, and a higher risk of seizure generation. These results provide valuable insights into the dynamics between amyloid pathology, network hyperactivity, and epileptogenesis in J20 mice.

### 3.5 Integrated Temporal Profile of Epileptogenesis and Amyloid Pathology

The longitudinal investigation into epileptogenic processes in AD highlights a staged progression, emphasizing the complex interplay between neural circuit dysfunction and amyloid pathology. To summarize the longitudinal changes across key hallmarks, the progression was categorized into three distinct phases:

1. At 9–15 weeks (Early Phase): Tg mice exhibited consistently elevated IED rates compared to nTg littermates, reflecting heightened neural excitability driven by APP overexpression. As memory impairments began to manifest during this period, the increasing ratio of CaMK2+ to PV+ neurons in Tg mice suggested a deteriorating excitatory-inhibitory imbalance. Concurrent iEEG recordings revealed a pronounced rise in patho-ripples and fast ripples, indicating the establishment of an epileptogenic network. Peaks in mossy fiber sprouting and unexpected mortality also occurred during this stage, highlighting the heightened vulnerability of hippocampal circuits to early pathological changes.
2. 15–21 weeks (Middle Phase): Amyloid plaque deposition began to develop, coinciding with the maturation of epileptogenic circuitry. This phase was characterized by a sustained elevation in patho-ripples and fast ripples, alongside an increase in HFO band power, reflecting enhanced neuronal excitability. Notably, these findings indicate that significant network dysfunction and epileptic activity precede the full deposition of amyloid plaques, suggesting a potential mechanistic link between early neural hyperactivity and amyloid pathology.
3. 21–27 weeks (Late Phase): Pathological electrophysiological markers such as patho-ripples and fast ripples plateaued, while amyloid plaque deposition being the sole biomarker continuing to rise. This upward trend in amyloid plaque accumulation persisted beyond the 27-week observation window.

These findings highlight a critical temporal relationship between neural circuit dysfunction and amyloid pathology, with early network hyperactivity and excitatory-inhibitory imbalance preceding the accumulation of amyloid plaques. By providing a detailed timeline of epileptogenesis in the context of AD, this study underscores the importance of early interventions targeting network hyperactivity to potentially mitigate downstream amyloid pathology and cognitive decline.

## 4. Discussion

Our longitudinal analysis reveals that epileptogenesis is not a downstream byproduct of AD pathology but a distinct and early component of disease progression. We demonstrate that pathological HFOs, including fast ripples and patho-ripples, emerge well before detectable amyloid plaque deposition and coincide with a progressive excitatory-inhibitory imbalance driven by early PV interneuron loss. There findings shift the prevailing view of hyperexcitability in AD and highlight neuronal network dysfunction as a mechanistic contributor-rather that a secondary consequence-of amyloid pathology.

### 4.1 Early Network Hyperactivity Precedes Amyloid Deposition

Hyperexcitability is increasingly recognized in both familial and sporadic AD, where network dysregulation rather than overt seizure activity appears an early pathogenic feature. Clinical studies demonstrate that subclinical epileptiform activity occur disproportionately in AD and closely track with cognitive decline[3, 4]. Our findings reinforce this view by demonstrating that pathological HFOs and the accompanying inhibitory interneuron deficits emerge well before detectable amyloid plaques. These results suggest that hyperexcitability functions as an upstream pathological event rather than a secondary consequence of late-stage neurodegeneration.

A central unresolved question is whether hyperexcitability actively contributes to amyloid deposition or reflects the downstream effects of APP overexpression. Although early-onset hyperexcitability has been consistently reported in many amyloid-based AD mouse models[11, 29], pinpointing its developmental origin is challenging because neonatal electrophysiology is rarely examined. In our study, J20 mice exhibited convulsive seizures (supplementary video) as early as 2 weeks of age, indicating an intrinsic excitatory bias long before plaque formation. This aligns with constitutive APP overexpression. Although APP normally contributes to neuronal growth, synaptic plasticity, and neuroprotection[30, 31], excessive APP enhances amyloidogenic processing and increases the production of soluble Aβ species. Soluble Aβ has been shown to disrupt synaptic transmission and promote network hyperactivity[32], whereas overt amyloid plaques only appear after these early circuit abnormalities are already established. Supporting this view, a recent study showed that elevated BACE1, the enzyme required for amyloidogenic APP cleavage, can cleave GABAAR subunit in APP23 mice and thereby promote hyperexcitability[33]. Paradoxically, BACE1 deletion and inhibition result in seizure phenotype[34], indicating that the relationship between APP processing and network excitability is more complex than a simple linear mechanism. Additional insights come from in vivo Ca²⁺ imaging studies. Hyperactive neuronal clusters have been detected near amyloid plaques[35], and hippocampal hyperactivity has been shown to precede plaque deposition and can be reduced by lowering Aβ production[36]. Together, these findings raise the possibility that amyloidogenic processing initiates epileptogenesis, not only amyloid pathology.

### 4.2 Inhibitory Network Destabilization Drives Early Hyperexcitability

Our findings reveal that PV interneuron loss emerges early in J20 mice and precedes amyloid plaque deposition, indicating that inhibitory failure is also a driver of early network hyperexcitability. PV interneurons provide fast perisomatic inhibition and coordinate pyramidal cell firing and maintain network oscillations, which are essential for cognitive processes[37]. Early vulnerability of PV interneurons offers a mechanistic explanation for the increasing excitability observed in the pre-plaque stage. This interpretation is supported by extensive evidence showing that impairments in PV interneuron function lead to memory deficits [38–40], and that Aβ-related toxicity contribute to PV interneuron loss[41–43]. Notably, restoring PV interneuron activity in APP/PS1 mice reduces amyloid burden and improves cognitive outcomes[28], suggesting that PV interneurons do not simply buffer excitability but actively influence the process of amyloid pathology. NPY upregulation is a well-characterized compensatory response to excessive excitatory drive in focal epilepsies, particularly in temporal lobe epilepsy, where it suppresses glutamatergic transmission and reduces seizure propensity[27]. The early surge of NPY in J20 mice likely reflects an attempt to stabilize hyperexcitable circuits during the initial phase of APP-driven imbalance, whereas its subsequent decline suggests that this compensatory mechanism becomes inadequate once inhibitory networks deteriorate. The combined progression of PV interneuron loss and the transient NPY response highlights a broader pattern of inhibitory system destabilization. As inhibitory control weakens, the resulting increase in HFO band power becomes a sensitive readout of emerging circuit instability, which forms the basis for the analysis in the following section.

### 4.3 Fast Ripples Reveal Pre-Amyloid Circuit Dysfunction

Pathological HFOs, particularly fast ripples, are well-established biomarkers for identifying epileptogenic zones and predicting seizure occurrence in epilepsy[22–24, 47, 48]. In chronic epilepsy models, increases in fast ripple frequency are believed to arise from interneuron loss and the strengthening of recurrent excitatory connections among pyramidal cells during latent epileptogenesis[44], a mechanism that aligns closely with the inhibitory deficits identified in our J20 mice. Only recently have HFOs been examined in the context of AD. The first report of HFOs in AD model was described by Gurevicius et al., who observed spontaneous HFOs in 4-month-old APPswe/PS1dE9 mice, although the oscillations occurred below the conventional pathological threshold for fast ripples[49]. Subsequent work by Lisgaras and Scharfman identified fast ripples in three distinct AD mouse models, supporting the idea that fast ripples may serve as electrophysiological indicators of AD-related circuit dysfunction[9]. Recently, Vossel et al. reported increased ripple and fast ripple activity in AD patients, providing strong clinical evidence that fast ripples may extend beyond classical epilepsy and serve as relevant biomarkers in human AD[50]. However, prior studies did not determine when pathological HFOs first appear. Animals were typically evaluated only after substantial amyloid deposition, leaving unresolved whether HFO abnormalities emerge before or only after plaque formation. Our longitudinal analysis fills this gap by demonstrating that patho-ripples and fast ripples escalate during the pre-plaque stage, when inhibitory network remodeling predominates, and well before amyloid plaques are detectable.

### 4.4 Temporal Biomarkers to Inform Early Intervention Strategies

The development of disease-modifying therapies in AD has centered primarily on targeting hallmark pathologies such as Aβ and tau. However, increasing evidence suggests that circuit-level dysfunction and hyperexcitability evolve alongside these molecular processes and may provide complementary information about early stages of disease progression. Within this framework, our longitudinal analysis offers a quantitative approach for situating electrophysiological biomarkers relative to amyloid pathology. By applying the Hill equation to biomarker trajectories in Tg and nTg mice, we were able to standardize developmental differences between groups and visualize the evolution of electrophysiological and histopathological changes across time. The resulting sigmoid (S-curve) fits, with R-squared values frequently exceeding 0.8, revealed that pathological electrophysiological patterns such as patho-ripples and fast ripples rise sharply before the appearance of detectable Aβ plaque in the hippocampus.

The S-curve model further delineated a critical period between weeks 15 and 21, a phase marked by maturation of excitatory-inhibitory balance, the development of pathological HFOs, and the initial emergence of amyloid plaques. This convergence of electrophysiological, cellular, and histopathological changes defines a narrow but actionable window in which therapeutic strategies that target network hyperexcitability may have maximal impact. Although anti-seizure clinical trials in AD have shown limited success[13], these studies predominantly enrolled patients with established dementia, likely beyond the optimal stage for neuromodulatory intervention. Our findings suggest that abnormalities in HFOs arise earlier, during a phase when network alterations may still be evolving. The S-curve framework may therefore help characterize the temporal relationship among electrophysiological markers, inhibitory circuit remodeling, and amyloid pathology. Such information may air future efforts to evaluate whether interventions that target hyperexcitability can influence downstream disease progresses. Overall, the present results support the value of using electrophysiological biomarkers to identify early circuit changes and motivate further studies aimed at determining how early modulation of network activity might impact the course of AD.

### 4.5 Study limitation and future directions

While our study provides novel insights into the temporal progression of epileptogenesis in AD, several limitations must be addressed: First, the hAPP-J20 model involves early and generalized APP overexpression, which may not fully reflect the heterogeneity of sporadic AD. Although HFOs have also been identified in AD patients, further validation across additional AD models and larger human cohorts will be necessary. Second, although early hyperexcitability clearly precedes amyloid deposition in our study, the causal relationship between these processes remains unresolved. Future work using more selective circuit-level or molecular manipulation will be needed to determine whether epileptiform activity contributes to amyloid accumulation or arises secondarily from soluble Aβ. Finally, although we observed early network dysfunction, our study did not include longitudinal behavioral assessment. Establishing clearer links between electrophysiological abnormalities and cognitive decline in human cohorts will be important for evaluating the translational significance of HFOs as early biomarkers.

## 5. Conclusion

Our study identifies a clear temporal sequence in which early network hyperactivity and excitatory-inhibitory imbalance emerge before amyloid plaque deposition in hAPP-J20 mice. This progression provides mechanistic insights into how circuit instability develops alongside amyloid pathology and highlights pathological HFOs as sensitive indicators of early network alterations. By delineating the dynamic evolution of epileptogenic circuits in AD, this study lays the groundwork for precision therapeutic strategies to modulate neural excitability and potentially slow disease progression.

## Supporting information

J20 mice exhibited convulsive seizures as early as 2 weeks of age postnatal.

## Abbreviation summary

AD: Alzheimer’s disease
Aβ: amyloid-beta
IEDs: interictal epileptiform discharges
HFOs: high-frequency oscillations
MCI: mild cognitive impairment
PCR: polymerase chain reaction
Tg: Transgenic
nTg: non-transgenic
iEEG: intracranial EEG
NREM: non-rapid eye movement
EA: epileptic activities
RMS: root mean square
SLL: short line length
NPY: neuropeptide Y
PV: Parvalbumin
ANOVA: analysis of variance
CNS: central nervous system
TLE: temporal lobe epilepsy
E/I: excitatory-inhibitory
DS: Down syndrome
S-curve: sigmoid curve

## Reference

1. Vossel KA, Ranasinghe KG, Beagle AJ, Mizuiri D, Honma SM, Dowling AF, Darwish SM, Van Berlo V, Barnes DE, Mantle M (2016) Incidence and impact of subclinical epileptiform activity in Alzheimer’s disease. Ann Neurol 80:858–870

2. Kamondi A, Grigg-Damberger M, Löscher W, Tanila H, Horvath AA (2024) Epilepsy and epileptiform activity in late-onset Alzheimer disease: clinical and pathophysiological advances, gaps and conundrums. Nat Rev Neurol 20:162–182

3. Ballerini A, Biagioli N, Carbone C, Chiari A, Tondelli M, Vinceti G, Bedin R, Malagoli M, Genovese M, Scolastico S (2025) Late-onset temporal lobe epilepsy: insights from brain atrophy and Alzheimer’s disease biomarkers. Brain 148:185–198

4. Vossel KA, Beagle AJ, Rabinovici GD, Shu H, Lee SE, Naasan G, Hegde M, Cornes SB, Henry ML, Nelson AB, Seeley WW, Geschwind MD, Gorno-Tempini ML, Shih T, Kirsch HE, Garcia PA, Miller BL, Mucke L (2013) Seizures and Epileptiform Activity in the Early Stages of Alzheimer Disease. JAMA Neurol 70:1158–1166. 10.1001/jamaneurol.2013.136

5. Vossel KA, Tartaglia MC, Nygaard HB, Zeman AZ, Miller BL (2017) Epileptic activity in Alzheimer’s disease: causes and clinical relevance. Lancet Neurol 16:311–322. 10.1016/S1474-4422(17)30044-3

6. Buzsáki G, Horváth Z, Urioste R, Hetke J, Wise K (1992) High-frequency network oscillation in the hippocampus. Science (80- ) 256:1025–1027. 10.1126/science.1589772

7. Staba RJ (2012) Normal and Pathologic High-Frequency Oscillations. In: Noebels JL, Avoli M, Rogawski MA, Olsen RW, Delgado-Escueta A V (eds). Bethesda (MD)

8. Staba RJ (2010) Normal and pathologic high-frequency oscillations. Epilepsia 51:21

9. Lisgaras CP, Scharfman HE (2023) High-frequency oscillations (250–500 Hz) in animal models of Alzheimer’s disease and two animal models of epilepsy. Epilepsia 64:231–246

10. Jones EA, Gillespie AK, Yoon SY, Frank LM, Huang Y (2019) Early hippocampal sharp-wave ripple deficits predict later learning and memory impairments in an Alzheimer’s disease mouse model. Cell Rep 29:2123–2133

11. Kam K, Duffy Á M, Moretto J, LaFrancois JJ, Scharfman HE (2016) Interictal spikes during sleep are an early defect in the Tg2576 mouse model of β-amyloid neuropathology. Sci Rep 6:20119

12. Palop JJ, Chin J, Roberson ED, Wang J, Thwin MT, Bien-Ly N, Yoo J, Ho KO, Yu G-Q, Kreitzer A (2007) Aberrant excitatory neuronal activity and compensatory remodeling of inhibitory hippocampal circuits in mouse models of Alzheimer’s disease. Neuron 55:697–711

13. Vossel K, Ranasinghe KG, Beagle AJ, La A, Pook KA, Castro M, Mizuiri D, Honma SM, Venkateswaran N, Koestler M (2021) Effect of levetiracetam on cognition in patients with Alzheimer disease with and without epileptiform activity: a randomized clinical trial. JAMA Neurol 78:1345–1354

14. Mucke L, Masliah E, Yu G-Q, Mallory M, Rockenstein EM, Tatsuno G, Hu K, Kholodenko D, Johnson-Wood K, McConlogue L (2000) High-Level Neuronal Expression of Aβ_1–42_ in Wild-Type Human Amyloid Protein Precursor Transgenic Mice: Synaptotoxicity without Plaque Formation. J Neurosci 20:4050 LP – 4058. 10.1523/JNEUROSCI.20-11-04050.2000

15. Vorhees C V, Williams MT (2006) Morris water maze: procedures for assessing spatial and related forms of learning and memory. Nat Protoc 1:848–858. 10.1038/nprot.2006.116

16. Minecan D, Natarajan A, Marzec M, Malow B (2002) Relationship of Epileptic Seizures to Sleep Stage and Sleep Depth. Sleep 25:56–61. 10.1093/sleep/25.8.56

17. von Ellenrieder N, Frauscher B, Dubeau F, Gotman J (2016) Interaction with slow waves during sleep improves discrimination of physiologic and pathologic high-frequency oscillations (80–500 Hz). Epilepsia 57:869–878. 10.1111/epi.13380

18. Ewell LA, Liang L, Armstrong C, Soltész I, Leutgeb S, Leutgeb JK (2015) Brain State Is a Major Factor in Preseizure Hippocampal Network Activity and Influences Success of Seizure Intervention. J Neurosci 35:15635 LP – 15648. 10.1523/JNEUROSCI.5112-14.2015

19. Janca R, Jezdik P, Cmejla R, Tomasek M, Worrell GA, Stead M, Wagenaar J, Jefferys JGR, Krsek P, Komarek V, Jiruska P, Marusic P (2015) Detection of Interictal Epileptiform Discharges Using Signal Envelope Distribution Modelling: Application to Epileptic and Non-Epileptic Intracranial Recordings. Brain Topogr 28:172–183. 10.1007/s10548-014-0379-1

20. Gardner AB, Worrell GA, Marsh E, Dlugos D, Litt B (2007) Human and automated detection of high-frequency oscillations in clinical intracranial EEG recordings. Clin Neurophysiol 118:1134–1143. 10.1016/j.clinph.2006.12.019

21. Lévesque M, Bortel A, Gotman J, Avoli M (2011) High-frequency (80–500Hz) oscillations and epileptogenesis in temporal lobe epilepsy. Neurobiol Dis 42:231–241. 10.1016/j.nbd.2011.01.007

22. Jacobs J, LeVan P, Chander R, Hall J, Dubeau F, Gotman J (2008) Interictal high-frequency oscillations (80–500 Hz) are an indicator of seizure onset areas independent of spikes in the human epileptic brain. Epilepsia 49:1893–1907. 10.1111/j.1528-1167.2008.01656.x

23. Wang S, Wang IZ, Bulacio JC, Mosher JC, Gonzalez-Martinez J, Alexopoulos A V, Najm IM, So NK (2013) Ripple classification helps to localize the seizure-onset zone in neocortical epilepsy. Epilepsia 54:370–376. 10.1111/j.1528-1167.2012.03721.x

24. Crépon B, Navarro V, Hasboun D, Clemenceau S, Martinerie J, Baulac M, Adam C, Le Van Quyen M (2010) Mapping interictal oscillations greater than 200 Hz recorded with intracranial macroelectrodes in human epilepsy. Brain 133:33–45. 10.1093/brain/awp277

25. Hong S, Beja-Glasser VF, Nfonoyim BM, Frouin A, Li S, Ramakrishnan S, Merry KM, Shi Q, Rosenthal A, Barres BA (2016) Complement and microglia mediate early synapse loss in Alzheimer mouse models. Science (80- ) 352:712–716

26. Webster SJ, Bachstetter AD, Nelson PT, Schmitt FA, Van Eldik LJ (2014) Using mice to model Alzheimer’s dementia: an overview of the clinical disease and the preclinical behavioral changes in 10 mouse models. Front Genet 5:88

27. Wickham J, Ledri M, Bengzon J, Jespersen B, Pinborg LH, Englund E, Woldbye DPD, Andersson M, Kokaia M (2019) Inhibition of epileptiform activity by neuropeptide Y in brain tissue from drug-resistant temporal lobe epilepsy patients. Sci Rep 9:19393

28. Hijazi S, Heistek TS, Scheltens P, Neumann U, Shimshek DR, Mansvelder HD, Smit AB, van Kesteren RE (2020) Early restoration of parvalbumin interneuron activity prevents memory loss and network hyperexcitability in a mouse model of Alzheimer’s disease. Mol Psychiatry 25:3380–3398

29. Kazim SF, Chuang S-C, Zhao W, Wong RKS, Bianchi R, Iqbal K (2017) Early-onset network hyperexcitability in presymptomatic Alzheimer’s disease transgenic mice is suppressed by passive immunization with anti-human APP/Aβ antibody and by mGluR5 blockade. Front Aging Neurosci 9:71

30. Chen G, Xu T, Yan Y, Zhou Y, Jiang Y, Melcher K, Xu HE (2017) Amyloid beta: structure, biology and structure-based therapeutic development. Acta Pharmacol Sin 38:1205–1235

31. Wilkins HM, Swerdlow RH (2017) Amyloid precursor protein processing and bioenergetics. Brain Res Bull 133:71–79

32. Martinsson I, Quintino L, Garcia MG, Konings SC, Torres-Garcia L, Svanbergsson A, Stange O, England R, Deierborg T, Li J-Y (2022) Aβ/amyloid precursor protein-induced hyperexcitability and dysregulation of homeostatic synaptic plasticity in neuron models of Alzheimer’s disease. Front Aging Neurosci 14:946297

33. Bi D, Bao H, Yang X, Wu Z, Yang X, Xu G, Liu X, Wan Z, Liu J, He J (2025) BACE1-dependent cleavage of GABAA receptor contributes to neural hyperexcitability and disease progression in Alzheimer’s disease. Neuron

34. Hitt BD, Jaramillo TC, Chetkovich DM, Vassar R (2010) BACE1-/-mice exhibit seizure activity that does not correlate with sodium channel level or axonal localization. Mol Neurodegener 5:1–14

35. Busche MA, Eichhoff G, Adelsberger H, Abramowski D, Wiederhold K-H, Haass C, Staufenbiel M, Konnerth A, Garaschuk O (2008) Clusters of hyperactive neurons near amyloid plaques in a mouse model of Alzheimer’s disease. Science (80- ) 321:1686–1689

36. Busche MA, Chen X, Henning HA, Reichwald J, Staufenbiel M, Sakmann B, Konnerth A (2012) Critical role of soluble amyloid-β for early hippocampal hyperactivity in a mouse model of Alzheimer’s disease. Proc Natl Acad Sci 109:8740–8745

37. Hu H, Gan J, Jonas P (2014) Fast-spiking, parvalbumin+ GABAergic interneurons: From cellular design to microcircuit function. Science (80- ) 345:1255263

38. Chung H, Park K, Jang HJ, Kohl MM, Kwag J (2020) Dissociation of somatostatin and parvalbumin interneurons circuit dysfunctions underlying hippocampal theta and gamma oscillations impaired by amyloid β oligomers in vivo. Brain Struct Funct 225:935–954

39. Miao C, Cao Q, Moser M-B, Moser EI (2017) Parvalbumin and somatostatin interneurons control different space-coding networks in the medial entorhinal cortex. Cell 171:507–521

40. Amilhon B, Huh CYL, Manseau F, Ducharme G, Nichol H, Adamantidis A, Williams S (2015) Parvalbumin interneurons of hippocampus tune population activity at theta frequency. Neuron 86:1277–1289

41. Sanchez-Mejias E, Nuñez-Diaz C, Sanchez-Varo R, Gomez-Arboledas A, Garcia-Leon JA, Fernandez-Valenzuela JJ, Mejias-Ortega M, Trujillo-Estrada L, Baglietto-Vargas D, Moreno-Gonzalez I (2020) Distinct disease-sensitive GABAergic neurons in the perirhinal cortex of Alzheimer’s mice and patients. Brain Pathol 30:345–363

42. Verret L, Mann EO, Hang GB, Barth AMI, Cobos I, Ho K, Devidze N, Masliah E, Kreitzer AC, Mody I (2012) Inhibitory interneuron deficit links altered network activity and cognitive dysfunction in Alzheimer model. Cell 149:708–721

43. Park K, Lee J, Jang HJ, Richards BA, Kohl MM, Kwag J (2020) Optogenetic activation of parvalbumin and somatostatin interneurons selectively restores theta-nested gamma oscillations and oscillation-induced spike timing-dependent long-term potentiation impaired by amyloid β oligomers. BMC Biol 18:1–20

44. Al Harrach M, Benquet P, Wendling F (2021) Long term evolution of fast ripples during epileptogenesis. J Neural Eng 18:46027

45. Puri BK, Ho KW, Singh I (2001) Age of seizure onset in adults with Down’s syndrome. Int J Clin Pract 55:442–444

46. Aller-Alvarez JS, Menéndez-González M, Ribacoba-Montero R, Salvado M, Vega V, Suárez-Moro R, Sueiras M, Toledo M, Salas-Puig J, Á lvarez-Sabin J (2017) Epilepsia mioclónica en el síndrome de Down y en la enfermedad de Alzheimer. Neurologia 32:69–73

47. Li L, Patel M, Almajano J, Engel Jr J, Bragin A (2018) Extrahippocampal high-frequency oscillations during epileptogenesis. Epilepsia 59:e51–e55

48. Ibarz JM, Foffani G, Cid E, Inostroza M, De La Prida LM (2010) Emergent dynamics of fast ripples in the epileptic hippocampus. J Neurosci 30:16249–16261

49. Gurevicius K, Lipponen A, Tanila H (2013) Increased cortical and thalamic excitability in freely moving APPswe/PS1dE9 mice modeling epileptic activity associated with Alzheimer’s disease. Cereb Cortex 23:1148–1158

50. Shandilya MCV, Addo-Osafo K, Ranasinghe KG, Shamas M, Staba R, Nagarajan SS, Vossel K (2025) High-frequency oscillations in epileptic and non-epileptic Alzheimer’s disease patients and the differential effect of levetiracetam on the oscillations. Brain Commun 7:fcaf041

